# 3D-Beacons: Decreasing the gap between protein sequences and structures through a federated network of protein structure data resources

**DOI:** 10.1101/2022.08.01.501973

**Authors:** Mihaly Varadi, Sreenath Nair, Ian Sillitoe, Gerardo Tauriello, Stephen Anyango, Stefan Bienert, Clemente Borges, Mandar Deshpande, Tim Green, Demis Hassabis, Andras Hatos, Tamas Hegedus, Maarten L Hekkelman, Robbie Joosten, John Jumper, Agata Laydon, Dmitry Molodenskiy, Damiano Piovesan, Edoardo Salladini, Steven L. Salzberg, Markus J Sommer, Martin Steinegger, Erzsebet Suhajda, Dmitri Svergun, Luiggi Tenorio-Ku, Silvio Tosatto, Kathryn Tunyasuvunakool, Andrew Mark Waterhouse, Augustin Žídek, Torsten Schwede, Christine Orengo, Sameer Velankar

**Affiliations:** European Molecular Biology Laboratory, European Bioinformatics Institute, Hinxton, UK; UCL, Department of Structural and Molecular Biology, London, UK; University of Basel, Biozentrum, Basel, Switzerland; European Molecular Biology Laboratory, EMBL Hamburg, Hamburg, Germany; DeepMind, London, UK; University of Padova, Department of Biomedical Sciences, Padova, Italy; University of Lausanne, SIB, Lausanne, Switzerland; Semmelweis University, Department of Biophysics and Radiation Biology, Budapest, Hungary; Netherlands Cancer Institute, Amsterdam, The Netherlands; Johns Hopkins University, Biomedical Engineering, Baltimore, MD, USA; Seoul National University, School of Biology, Seoul, South Korea

**Author notes:** these authors contributed equally (Mihaly Varadi, Sreenath Nair, Ian Sillitoe, Gerardo Tauriello).

## Abstract

While scientists can often infer the biological function of proteins from their 3-dimensional quaternary structures, the gap between the number of known protein sequences and their experimentally determined structures keeps increasing. A potential solution to this problem is presented by ever more sophisticated computational protein modelling approaches. While often powerful on their own, most methods have strengths and weaknesses. Therefore, it benefits researchers to examine models from various model providers and perform comparative analysis to identify what models can best address their specific use cases. To make data from a large array of model providers more easily accessible to the broader scientific community, we established 3D-Beacons, a collaborative initiative to create a federated network with unified data access mechanisms. The 3D-Beacons Network allows researchers to collate coordinate files and metadata for experimentally determined and theoretical protein models from state-of-the-art and specialist model providers and also from the Protein Data Bank.

## Introduction

Proteins are essential building blocks of almost every biological process; therefore, understanding their functions is critical to many applications, from drug discovery^1,2^ to tackling environmental challenges such as plastic pollution^3^. Accurate information on the structure of a protein, especially in the context of its biological assembly, can help scientists understand and modulate its function^4,5^.

Unfortunately, gaining such insights regarding the function of proteins through their structures is severely hampered by the lack of high-quality, experimentally determined structures. As of 2022, the Universal Protein Resource (UniProt) contains around 204 million non-redundant amino acid sequences, while the Protein Data Bank (PDB)^6,7^ contains around 190,000 PDB entries mapped to approximately 52,000 UniProt accessions. In other words, less than 0.03% of all the known protein sequences have experimentally determined atomic resolution structures. As sequencing becomes more accessible, the gap between protein sequences and structures increases (Figure 1).

**Figure 1.**
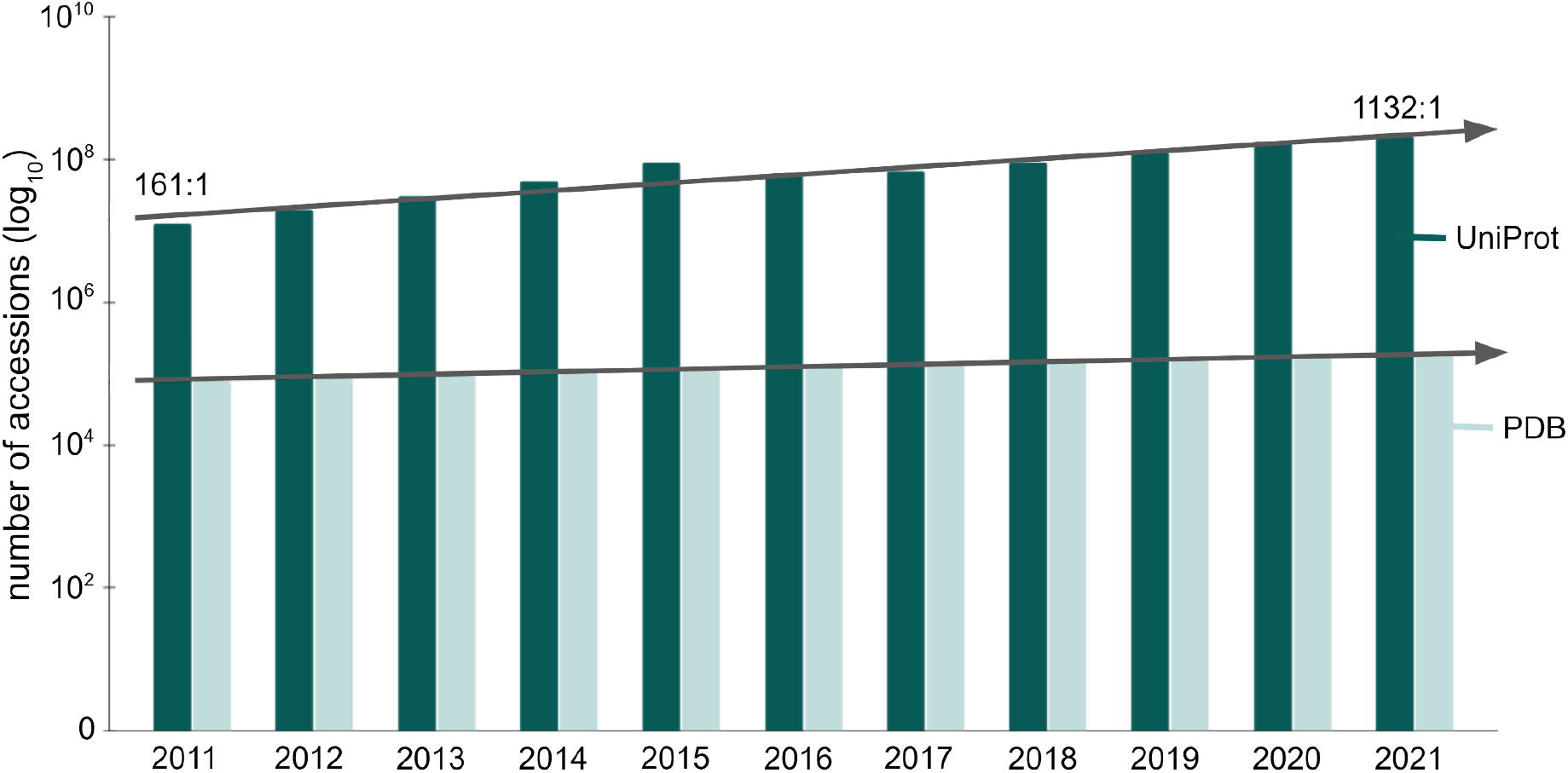
Growth of the UniProt and the PDB databases.

This figure shows the number of accessions (on a logarithmic scale) throughout the past decade. In 2011, the UniProt had 161× as many protein sequences as the number of PDB entries. This ratio grew by an order of magnitude, and was 1132 to 1 in 2021, showing that the gap between known protein sequences and their structures keeps increasing.

A practical approach to addressing this challenge relies on high-accuracy computational models to complement the experimentally determined structures when the latter are unavailable for a certain protein of interest^8^. The thermodynamic hypothesis postulates that within certain limitations, the native structure is determined only by the protein’s amino acid sequence^9^,^10^. Indeed, the past 50 years saw the development of many algorithms and scientific software to predict protein structures^11^,^12^. An approach developed early in this field was to use homologous protein structures as templates. Several modelling tools and data resources have long provided access to such models, for example, the SWISS-MODEL and the ModBase web-services and databases^13–15^. In 2021 the field saw tremendous advances with tools such as AlphaFold and RoseTTAFold achieving much higher accuracy for “*de novo*” predictions without homologous templates than ever before^16,17^. This new generation of prediction tools makes it possible to try and predict the structure of virtually any known protein based on its sequence.

While these new techniques are increasingly accurate, it is important that they are supplemented with reliable estimates of model confidence both for the whole model and locally for each residue. Researchers should not expect all predictions to be equally accurate neither globally nor in every region, and confidence estimates should hence be used to determine if a predicted structure can be used for downstream analysis^18^. Commonly used model confidence methods aim to predict the global and local similarity of the model compared to the correct coordinates if those coordinates were provided by an experimentally determined structure. In recent years, several model prediction methods such as SWISS-MODEL^14^, RoseTTAFold^17^ and AlphaFold^16^ have chosen the superposition-free lDDT score^19^ (pLDDT for AlphaFold^16^) as a similarity metric to provide model confidence for their own models. The lDDT score measures differences in interatomic distances within a short radius between model and reference structure. It has been shown that superposition-free measures are robust with respect to domain movements and have advantages for the analysis of local structural details^20^. Similarly, superposition-free measures have been used for a long time in the creation of experimental structure models^21^.

Another important consideration when relying on any structure prediction tool is to consider its limitations. While structures in the PDB have the advantage of experimental data backing the coordinates, enabling experimental as well as geometric validation, it is a relatively small data set, as discussed above. Template-based models have the distinct advantage of enabling the mapping of a model to homologues with known structures, thus mapping to experimentally derived structures which can be in distinct conformational states or in complex with other molecules. Some tools excel at general-purpose protein structure modelling; others specialise in placing relevant ligands in the context of a model or representing conformational flexibility with ensembles of potential conformations^14,16,22–24^ (Figure 2). For example, AlphaFold 2.0 cannot perform docking of small molecules, even if they are obligate ligands of the proteins, such as Zinc-finger proteins. However, data resources such as AlphaFill can tackle this problem by building on existing models and adding known ligands to these structures^23^ (Figure 2A). On the other hand, the central repository of AlphaFold models, the AlphaFold Structure Database, only contains predictions for single polypeptide chains and not necessarily the functional forms of proteins^25^. In the case of multimeric complexes, the functional form can include several polypeptide chains. Since the number of known protein complexes is immense, having a comprehensive database for complex structures soon is rather unlikely. Therefore, integrating 3D data from experts in specialised fields of proteins is important, as demonstrated by physiologically and pathologically relevant transmembrane ABC half transporters^26^ and by a set of computed structures of core eukaryotic protein complexes deposited in the ModelArchive^27^. Databases such as the Small-Angle Scattering Biological Data Bank (SASBDB)^28^ and the Protein Ensemble Database (PED)^22^ highlight the dynamic nature of intrinsically disordered proteins (Figure 2B). Small-angle scattering provides low-resolution information on the shape and size of biological macromolecules in solution, but it also offers powerful means for the quantitative analysis of flexible systems, including intrinsically disordered proteins (IDPs)^29^. This data together with *ab initio* modelling approaches can be utilised to generate an experimentally validated pool of IDP models. PED provides access to such conformational ensembles, but also those based on other experimental approaches. Considering the limitations of certain tools highlights the importance of using models and methods from various synergistic software and data providers to mitigate the weaknesses of individual modelling techniques.

**Figure 2.**
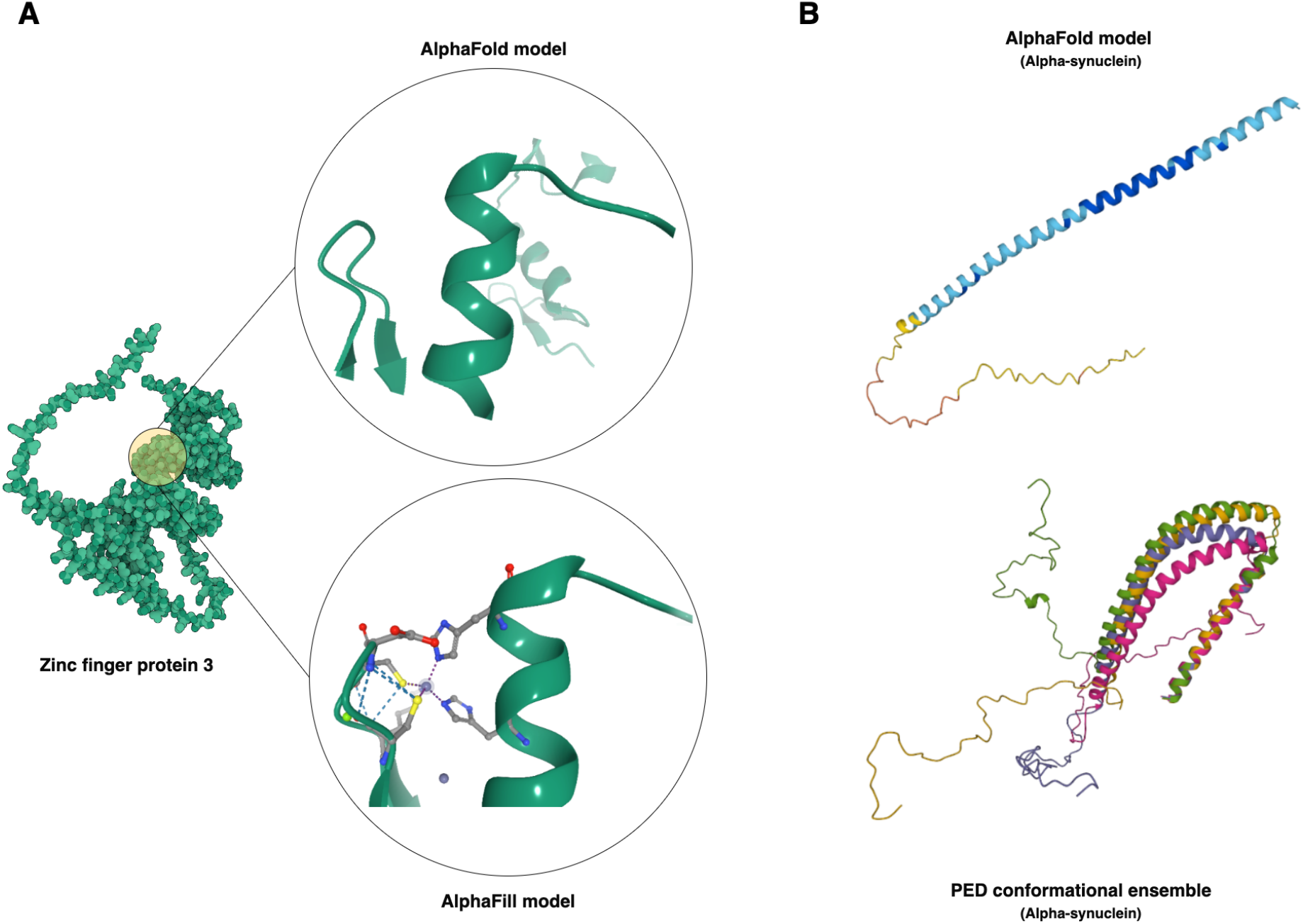
Highlighting the strengths and weaknesses of modelling techniques.

Each modelling approach has limitations and specific strengths. For example, AlphaFill complements AlphaFold models by placing obligate ligands in their contexts (panel A). Other data providers, such as the Protein Ensemble Database, provide conformational ensembles for intrinsically disordered proteins (IDPs), for example, for the human Alpha-synuclein (panel B).

While many prediction software and several publicly accessible data resources host and archive protein structures, these resources are fragmented and often rely on their own data standards to describe the necessary meta-information essential for providing context for a specific model. They also offer distinct data access mechanisms, requiring the users to learn multiple sets of technical details when interacting with various resources. The lack of standardisation can severely impede the comparative analysis of these models, making it difficult to gain valuable insights.

Here, we present the 3D-Beacons Network (https://3d-beacons.org), an open, collaborative platform for providing programmatic access to 3-dimensional coordinates and their standardised meta-information from both experimentally determined and computationally modelled protein structures.

## Results

The 3D-Beacons Network is an open collaboration between providers of experimentally determined and computationally predicted protein structures. To date, ten data providers make their protein structures available through this platform (Table 1). The consortium is guided by a collaboration agreement that prospective data providers agree to comply with (https://www.ebi.ac.uk/pdbe/pdbe-kb/3dbeacons/guidelines). Importantly, all the data provided through the network must be freely available for academic and commercial use under Creative Commons Attribution 4.0 (CC-BY 4.0, https://creativecommons.org/licenses/by/4.0/) licence terms.

**Table 1.** Members of the 3D-Beacons Network.

| Data provider | Model category | Number of structures* |
| --- | --- | --- |
| AlphaFill | Template-based | 995,411 |
| AlphaFold DB | Ab initio | 214,684,311 |
| Genome3D | Template-based | *In progress* |
| HegeLab | Ab initio | 15 |
| isoform.io | Ab initio | 48,551 |
| ModelArchive | Ab initio / template-based | 1,106 |
| PDBe | Experimentally determined | 190,639 |
| PED | Conformation ensembles | 275 |
| SASBDB | Experimentally determined | 3,912 |
| SWISS-MODEL Repository | Template-based | 2,216,915 |

\* Numbers are accurate as of 29th July 2022

The 3D-Beacons Network is based on an infrastructure that helps providers of protein structures to standardise their meta-information, and easily link their model files to a centralised search engine, called the 3D-Beacons Hub API (application programming interface) (Figure 3). Each data provider has its 3D-Beacon connected to the central Hub. The Hub is the public access point through which the users (or other data services) can retrieve models from any members. This allows users to get all structures for a given UniProt accession instead of manually retrieving them from all the different structure providers.

**Figure 3.**
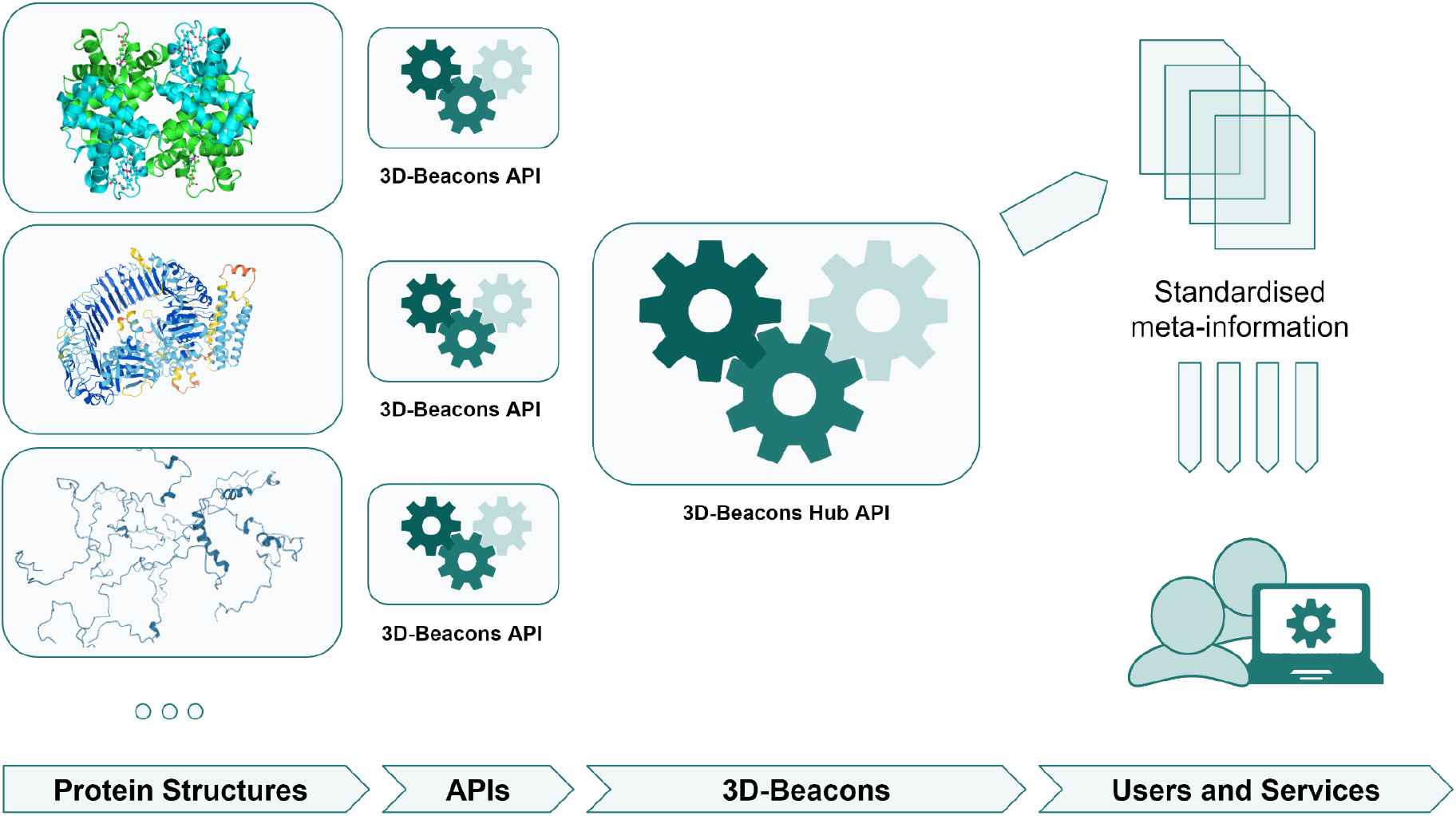
Schematic overview of the 3D-Beacons Network.

Thanks to the standardised data formats, the infrastructure assures complete transparency in data provenance and allows users to easily compare protein structures and their relevant meta-information. This initiative has evolved in parallel with efforts to improve the standardisation of the coordinate files for theoretical models. In particular, members of the 3D-Beacons Network contributed to the ModelCIF extension of the PDBx/mmCIF format, which supports more exhaustive meta-information and includes mappings to the corresponding UniProt accessions next to the atomic coordinates.

Data providers standardise their meta-information and make their models available through 3D-Beacons API instances. The 3D-Beacons Registry links these instances to the central 3D-Beacons Hub API, which can be openly accessed by the scientific community and other data services.

While the primary purpose of 3D-Beacons is to provide efficient and scalable programmatic access to protein structures, we also offer a graphical user interface that allows researchers to get an overview of the available protein structures. For example, users can view all the available data from any member data provider for the human Cellular tumour antigen p53 protein by searching based on its UniProt accession: https://www.ebi.ac.uk/pdbe/pdbe-kb/3dbeacons/search/P04637 (Figure 4).

**Figure 4.**
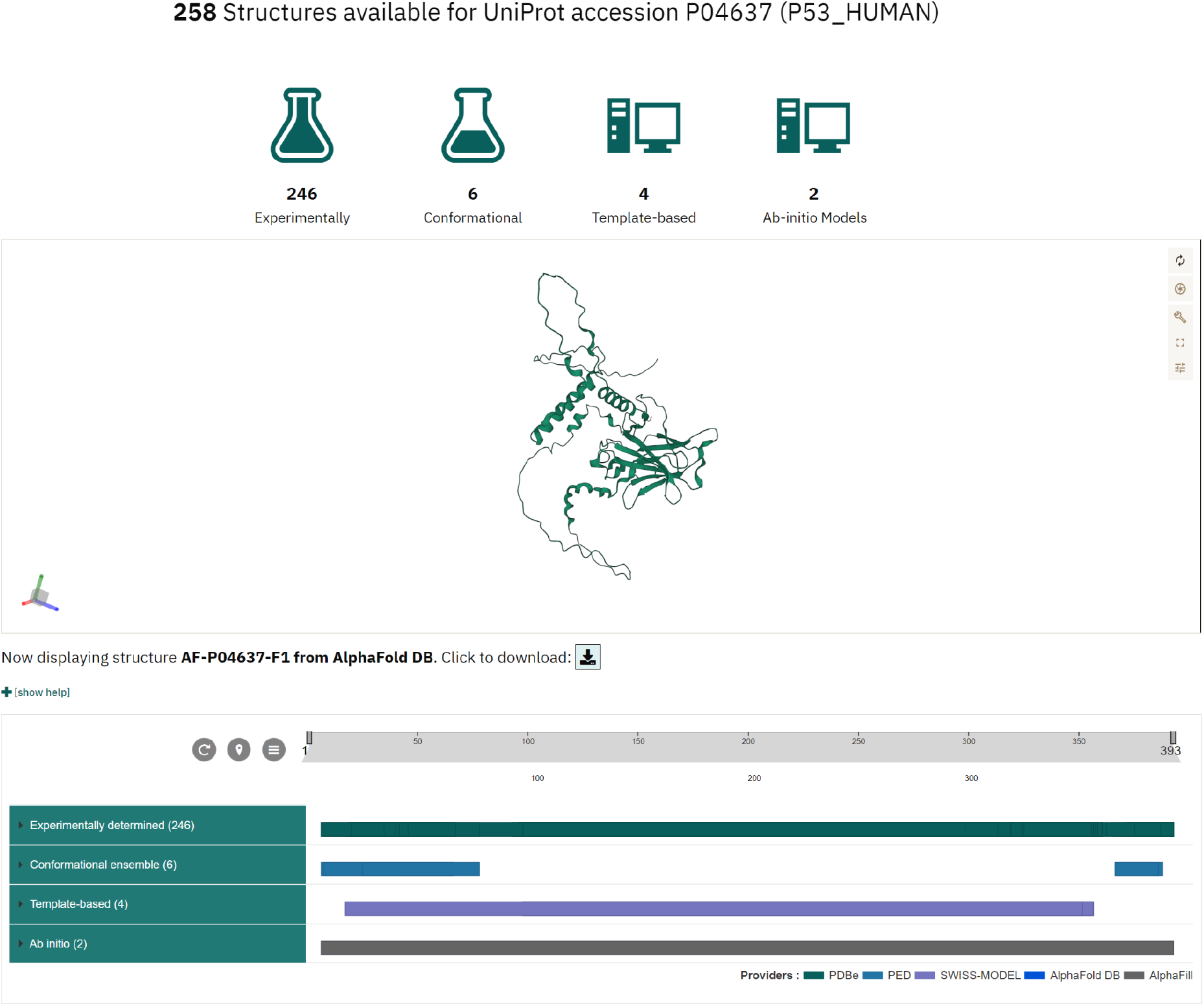
Graphical user interface of 3D-Beacons.

While the main focus of the 3D-Beacons Network is to provide programmatic access to experimentally determined and computationally predicted protein structures, we also provide a graphical user interface where researchers can query for specific proteins using UniProt accessions. This interface displays which section of the protein sequence the models cover and provides an interactive 3D view.

We divided the protein structures into four categories: 1) experimentally determined; 2) template-based; 3) *ab-initio*; and 4) conformational ensembles. We defined the categories as follows:

**Experimentally determined** structures are based on data from techniques such as X-ray crystallography, cryo-electron microscopy, nuclear magnetic resonance spectroscopy or small-angle scattering. This category is exemplified by structures in the PDB and the SASBDB databases.
**Template-based** models use alignments to similar sequences with known structure (i.e. templates) as their main input. SWISS-MODEL is an example of data providers with such models.
**Ab-initio** models can use templates as an auxiliary input, but do not depend on them. AlphaFold models are considered *ab-initio* in this framework.

Finally, **conformational ensembles** are created using a combination of experimental data and computational modelling, yielding a large number of possible conformations. Ensembles in the PED database are an example of this category.

Researchers can view the number of models under each category and inspect which parts of the amino acid sequences are covered by which models in a 2D viewer, PDB ProtVista^30^. Users can also display the structures using an embedded 3D molecular graphics viewer, Mol*^31^, and download the models in PDB or mmCIF formats.

## Discussion

The purpose of the 3D-Beacons Network is to standardise the representation of protein structure models and associated metadata and to provide efficient, high-throughput programmatic access to experimentally determined and theoretical models and their standardised metadata. The current version (as of 29th July 2022) of 3D-Beacons supports querying any number of UniProt accessions, while future updates are planned to collate models based on other identifiers such as taxonomy IDs or domain IDs. This platform enables both the scientific community and developers of data visualisation and data providing services to access and seamlessly integrate 3D models from various protein structure data providers.

While designing the data access points and data formats, we had extensive discussions with scientists and developers who provided specific use cases that are relevant to their work.

We used this data to drive the development of 3D-Beacons, starting with the most frequently requested data, i.e. information keyed on UniProt accessions, that can answer the question “What experimental or theoretical structures are available for my protein of interest?”. Going forward, we will address more of the collated use cases, such as searching by sequence or by gene identifiers and selecting structures based on protein families. Already, the API endpoints of 3D-Beacons provide easy access to models from sparse and fragmented data resources, supporting researchers and software developers alike.

For example, the 3D-Beacons infrastructure allows users of Jalview, a workbench for creating multiple sequence alignments (MSAs) and analysing them, to discover 3D models for MSAs of proteins from the UniProt and place them in the context of genetic variation from Ensembl^32^. It can also visualise local model quality scores such as pLDDT.

The Protein Data Bank in Europe – Knowledge Base (PDBe-KB)^33^ displays all the experimentally determined and computationally predicted structures for proteins of interest on their aggregated views of proteins. To retrieve metadata and the location of model files, it uses the 3D-Beacons Hub API. This integration also allows PDBe-KB to display functional and biophysical annotations both for theoretical models in addition to experimentally determined structures.

The SWISS-MODEL Repository (SMR)^13^ fetches models from AlphaFold DB and the ModelArchive using the 3D-Beacons Hub API. SMR displays these models next to homology models from SWISS-MODEL^14^ and experimental structures from the PDB^6^ to facilitate comparative analysis. SMR also takes advantage of the confidence measure information, and the models are displayed with a consistent colouring based on these confidence metrics.

## Methods

The infrastructure of the 3D-Beacons Network consists of a registry, a hub and the data access implementations. The 3D-Beacons Network is open to data providers of protein structures. Such data resources are invited to contact the 3D-Beacons consortium to discuss ways their data can be linked. Briefly, the common steps are as follows: Data providers review the consortium guidelines (https://3d-beacons.org/guidelines) and the latest API specification.The data providers then convert their metadata to the specified format and make these data available either through their APIs or by setting up a 3D-Beacon client. Once these steps are completed, the registry can be updated to link the new data resource with the 3D-Beacons Hub API. The following sections below give more detailed information on each of these elements of the infrastructure.

### 3D-Beacons Registry

The 3D-Beacons Registry is a transparent, publicly accessible registry that stores information on all the data providers linked to the 3D-Beacons Network. The registry is available on GitHub (https://github.com/3D-Beacons/3d-beacons-registry). It contains information on the public URLs of data providers, a brief description of the protein structures they provide, and a list of API endpoints they support. For example, PDBe^34^’^35^ supports the API endpoint that is keyed on a UniProt accession, and that provides high-level information about the models, while SMR^13^ supports both the high-level and the detailed API endpoints, which additionally provides per-chain and per-residue information on the models.

### 3D-Beacons data exchange format

The API endpoints comply with the data exchange format, which the 3D-Beacons members collaboratively design and improve. We defined the data exchange format as a JavaScript Object Notation (JSON) specification, an industry-standard format for sharing textual meta-information. The specification is available on Apiary (https://3dbeacons.docs.apiary.io/#) and GitHub (https://github.com/3D-Beacons/3d-beacons-specifications/releases).

### 3D-Beacons client

Members of the 3D-Beacons Network can either implement their own API endpoints according to the API specification described above, or they can install a local instance of the 3D-Beacons client. This client is a Docker-containerized, lightweight Python package that can import and parse PDB or mmCIF formatted protein structure files and their corresponding meta-information (in JSON format). It also includes capabilities to add model confidence scores using QMEANDisCo^36^ if models do not already include comparable scores such as pLDDT. QMEANDisCo, which is used internally by SWISS-MODEL, can be applied to models from any provider and has proven to be an accurate confidence predictor for homology modelling and some *ab initio* methods^20^. The client indexes the collated data in an embedded MongoDB database instance and exposes the information through an embedded API implementation that complies with the 3D-Beacons API specifications. The client is freely available on GitHub (https://github.com/3D-Beacons/3d-beacons-client).

### 3D-Beacons Hub API

At the core of the 3D-Beacons infrastructure lies the Hub API, a programmatic aggregator of the meta-information from all the member data providers. We implemented the Hub API using the FastAPI framework. The source code of this API is available at https://github.com/3D-Beacons/3d-beacons-hub-api. This API relies on the previously described registry to retrieve information on which data provider supports which specific API endpoints. It aggregates data and provides its own API endpoints that researchers, services and software can directly access to retrieve the location of available model files and their corresponding meta-information, such as the overall model quality or residue-level confidence measures.

### 3D-Beacons front-end

Finally, we provide a graphical user interface that contains documentation and showcases the information one can retrieve using the 3D-Beacons Hub API at https://3d-beacons.org. We implemented this interface using the Angular framework, and it relies on the sequence feature viewer, PDB ProtVista^30^ and the 3D molecular graphics viewer, Mol*^31^. The source code of this front-end application is available at https://github.com/3D-Beacons/3d-beacons-front-end.

## Acknowledgements

Work on creating the 3D-Beacons infrastructure was primarily funded by the BBSRC grant BB/S020071/1. We would like to acknowledge the contribution of ELIXIR BioHackathon participants in 2020 and 2021 who helped improve various aspects of the 3D-Beacons infrastructure. G.T., A.M.W., S.B. and T.S. acknowledge funding from ELIXIR and the SIB Swiss Institute of Bioinformatics. D.I.S. and D.S.M. acknowledge funding from the German Ministry of Science and Education Grant Number: 16QK10A-SAS-BSOFT. M.S. acknowledges support from the National Research Foundation of Korea (NRF) Grant Number: 2021-R1C1-C102065.

## Author contributions

M.V. created the initial draft, handled project and data management at PDBe and AlphaFold DB, and designed and developed the 3D-Beacons web pages. S.N. led the development of the 3D-Beacons registry, Hub API and PDBe and AlphaFold API implementation, as well as contributed to the development of the 3D-Beacons client and the web pages. I.S. worked on the 3D-Beacons client, and on the Genome3D beacon. G.T. contributed to the management of 3D-Beacons. M.V., G.T., A.M.W., S.B. contributed to the API design. A.L., N.A. provided management support for AlphaFold DB. A.H., S.T., L.T-K., E.S., D.P. contributed to the PED beacon. A.M.W., S.B., G.T. contributed to the SWISS-MODEL and ModelArchive beacons. S.B. contributed QMEANDisCo for the client. S.B., G.T. connected with the ModelCIF working group. T.H., E.S. contributed to the HegeLab beacon. M.L.H., R.P.J. contributed to the AlphaFill beacon. C.B., Dm.S, D.M. contributed to the SASBDB beacon. M.S., M.J.S., S.L.S. contributed to the isoform.io beacon. M.D., S.A. worked on data visualisation and infrastructure. S.V., C.O., T.S. provided oversight as co-PIs. I.S., G.T., D.M., T.H., S.V., R.J., E.S. contributed to the manuscript drafts. Every co-author reviewed the final manuscript.

